# Cytokine and chemokine profile in patients hospitalized with COVID-19: A comparative study

**DOI:** 10.1101/2022.03.17.484837

**Authors:** Abdisa Tufa, Tewodros Haile Gebremariam, Tsegahun Manyazewal, Tewodros Getinet, Dominic-Luc Webb, Per M. Hellström, Solomon Genet

## Abstract

Abnormal cytokine and chemokine concentrations during SARS-CoV-2 infection may represent disease severity. We aimed to assess plasma cytokine and chemokine concentrations in patients with SARS-CoV-2 in Addis Ababa, Ethiopia. In this study, 260 adults: 126 hospitalized patients with confirmed COVID-19 sorted into severity groups: severe (n=68) and mild or moderate (n=58), and 134 healthy controls were enrolled. We quantified 39 plasma cytokines and chemokines using multiplex ELISA. Spearman rank correlation and Mann-Whitney U test were used to identify mechanistically coupled cytokines/chemokines and compare disease severity. Compared to healthy controls, patients with COVID-19 had significantly higher levels of interleukins 1α, 2, 6, 7, 8, 10 and 15, C-reactive protein (CRP), serum amyloid A (SAA), intercellular adhesion molecule 1 (ICAM-1), vascular cell adhesion protein 1 (VCAM-1), IFN-γ-inducible protein-10 (IP-10), macrophage inflammatory protein-1 alpha (MIP-1α), eotaxin-3, interferon-gamma (IFN-ϒ), tumor necrosis factor-α (TNF-α), basic fibroblast growth factor (bFGF), placental growth factor (PlGF), and fms-like tyrosine kinase 1 (Flt-1). Patients with severe COVID-19 had higher IL-10 and lower macrophage-derived chemokine (MDC) compared to the mild or moderate group (*P<0.05*). In the receiver operating characteristic curve, SAA, IL-6 and CRP showed strong sensitivity and specificity predicting the severity and prognosis of COVID-19. Greater age and higher CRP had a significant association with disease severity (*P<0.05*). Our findings reveal that CRP, SAA, VCAM-1, IP-10, MDC and IL-10 levels are promising biomarkers for COVID-19 disease severity, suggesting that plasma cytokines/chemokines could be used as warning indicators of COVID-19 severity, aid in COVID-19 prognosis and treatment.

**IMPORTANCE:** SARS-CoV-2 triggers inflammatory reaction resulting in respiratory discomfort and in critical case may result in death. Cytokines and chemokines are inflammatory biomarkers that regulate and determine the nature of immune responses. Measuring cytokine and chemokine levels is useful in stratification, management and treatment of COVID-19 patients as well as guide resource allocations and therapeutic options. Here, we examined ctytokine and chemokine profiles in COVID-19 patients. Understanding how distinct cytokines and chemokines change over time as COVID-19 disease progresses might aid clinicians in detecting severe illness earlier and thereby improve patient prognosis.

## INTRODUCTION

SARS-CoV-2 has unusually potent transmission and toxicity. It shares severe flu-like symptoms and acute respiratory distress syndrome of zoonotic diseases with other SARS variations (e.g., SARS-CoV) and the Middle East Respiratory Syndrome (MERS) (1). Individuals who test positive for SARS-CoV-2 by molecular diagnostics, such as reverse-transcriptase polymerase chain reaction (RT-PCR), are initially classified as asymptomatic or pre-symptomatic. Furthermore, COVID-19 symptomatology is commonly related to fever, cough, dyspnea and myalgia or weariness (2). Sputum production, headaches, hemoptysis and diarrhea are all minor symptoms. Infectious pneumonia is the hallmark of severe disease and consequences may include acute respiratory distress syndrome (ARDS), abrupt heart damage and secondary infection (3). The severity of a disease is determined by its symptomatology. The mean incubation period for COVID-19 was 6 days globally, but around 7 days on the Chinese mainland, which will aid in determining the time of infection and making disease control decisions (4).

Many investigations have indicated that COVID-19 patients had increased levels of circulating proinflammatory cytokines (reviewed in 5). During the rapid course of COVID-19, a cytokine storm sometimes ensues, altering the immune system by lowering lymphocyte counts, particularly T cells (6). When the immune system as a whole is disrupted, immune cells release a huge number of pro-inflammatory cytokines and chemokines, which exacerbates the cytokine storm (7). In the most severe cases of COVID-19, IL-6 levels are greatly elevated and this is one of the factors that leads to cytokine production (8). In COVID-19 patients, viral-induced hyper-inflammation is strongly linked to disease severity and it is even the leading cause of mortality (9). As a result of an erroneous immune response, COVID-19’s critical and severe cases are characterized by sepsis and multiple organ failure (10).

Chemokines are important inflammatory mediators that help the immune system respond to infections; nonetheless, excessive production is the major cause of hyperinflammation (11). According to a meta-analysis, chemokines (MCP-1, IP-10 and Eotaxin) were shown to be higher in severe COVID-19 patients than in moderate cases (12). IP-10 and MCP-1 are biomarkers that have been associated to the severity of COVID-19 disease and the risk of death in COVID-19 patients (13). COVID-19 cases progressively increased in Ethiopia, with a total of 467,288 new cases and 7,417 deaths as of February 12^th^, 2022.

The objective of this study was to assess circulating concentrations of cytokines and chemokines in COVID-19 patients in Addis Ababa, Ethiopia, in order to investigate their diagnostic relevance, prognosis and classification of disease severity in separating COVID-19 and healthy controls group.

## RESULTS

### Demography and clinical characteristic

There were 126 COVID-19 patients and 134 healthy control subjects. Males made up 155 (60 %) of participants. Patients were separated into two groups: severe (68, 54 %) and mild or moderate (58, 46 %). The median age of mild or moderate (32 years of age, IQR 20-78) versus severe (60 years of age, IQR 22-86) groups was significant (Table 1). There were 64 COVID-19 patients (50.8 %) with either single or multiple morbidities such as diabetes (16 %), hypertension (14 %), cardiovascular disease (CVD) (17 %), cancer (17 %) and chronic lung disease (10 %). In the severe group, the proportion of diabetes and chronic lung disease were considerably higher than in the mild or moderate group (Table 2).

**TABLE 1.**
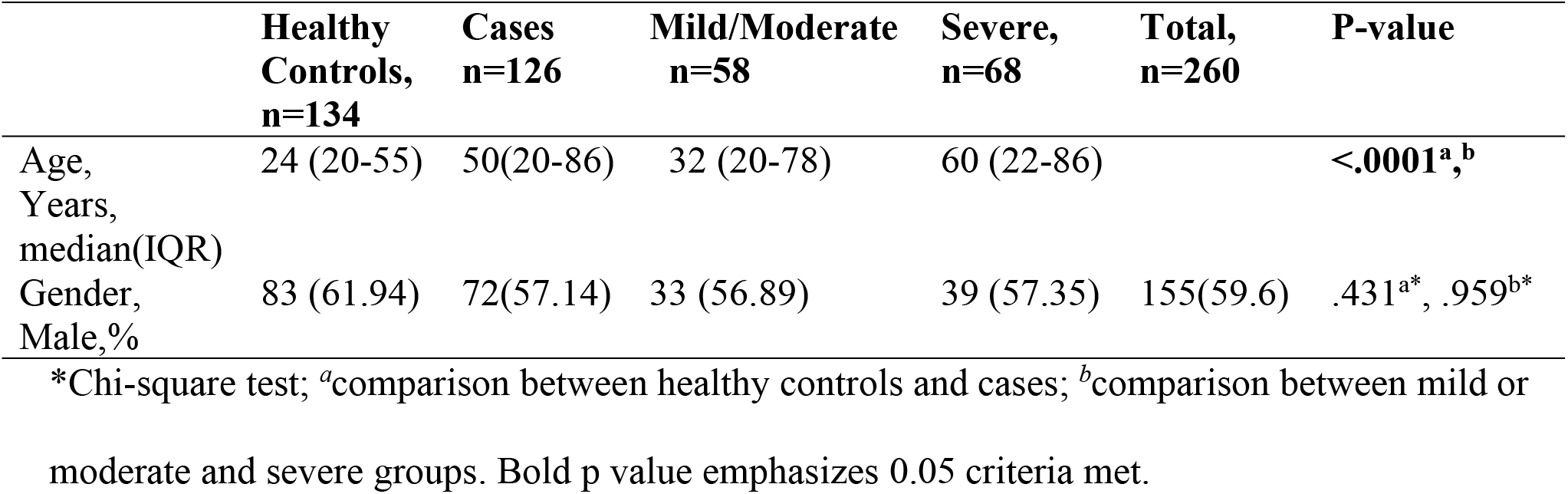
Demographics characteristic of COVID-19 patients and healthy controls

|  | Healthy Controls, n=134 | Cases n=126 | Mild/Moderate n=58 | Severe, n=68 | Total, n=260 | P-value |
| --- | --- | --- | --- | --- | --- | --- |
| Age, Years, median(IQR) | 24 (20-55) | 50(20-86) | 32 (20-78) | 60 (22-86) |  | <.0001<sup>a, b</sup> |
| Gender, Male, % | 83 (61.94) | 72(57.14) | 33 (56.89) | 39 (57.35) | 155(59.6) | .431<sup>a*</sup>, .959<sup>b*</sup> |

\*Chi-square test; <sup>a</sup>comparison between healthy controls and cases; <sup>b</sup>comparison between mild or moderate and severe groups. Bold p value emphasizes 0.05 criteria met.

**TABLE 2.** Comorbidity by COVID severity classification

|  | All patients n (%) | Mild/Moderate n (%) | Severe n (%) | P-value |
| --- | --- | --- | --- | --- |
| **Any of following** |  |  |  |  |
| Diabetes | 20 (15.9) | 5 (8.6) | 15 (22.1) | **.040** |
| Hypertension | 17 (13.5) | 5 (8.6) | 12 (17.6) | .139 |
| CVD | 21 (16.7) | 10 (17.2) | 11 (16.2) | .873 |
| Cancer | 21 (16.7) | 12 (20.7) | 9 (13.2) | .263 |
| Chronic lung disease | 12 (9.5) | 2 (3.4) | 10 (14.7) | **.032** |
| Comorbidity | 64 (50.8) | 27 (46.6) | 37 (54.4) |  |
| Non-comorbidity | 62 (49.2) | 31 (53.4) | 31 (45.6) | .379 |
| **Total patients (n)** | 126 | 58 | 68 |  |

Chi-square test was used. Bold p value emphasizes 0.05 criteria met. CVD, Chronic cardiovascular disease. Individuals have multiple comorbidity illness.

### Cytokines and chemokines plasma concentrations

Circulating cytokine and chemokine concentrations are summarized in table 3. Circulating concentrations of 19 cytokines or chemokines (interleukins 1α, 2, 6, 7, 8, 10 and 15, CRP, SAA, ICAM-1, VCAM-1, IP-10, MIP-1α, Eotaxin-3, IFN-ϒ, TNF-α, bFGF, PlGF and Flt-1) in COVID-19 patients were increased compared to the healthy controls (P < 0.05,Table 3). Even after age was adjusted (comparing age ≤ 40 years in both COVID-19 patients and healthy control group), the majority of the median levels of cytokines and chemokines in the COVID-19 patients were greater than the healthy control group and statistically significant (P < 0.05, Table 5).

**TABLE 3.**
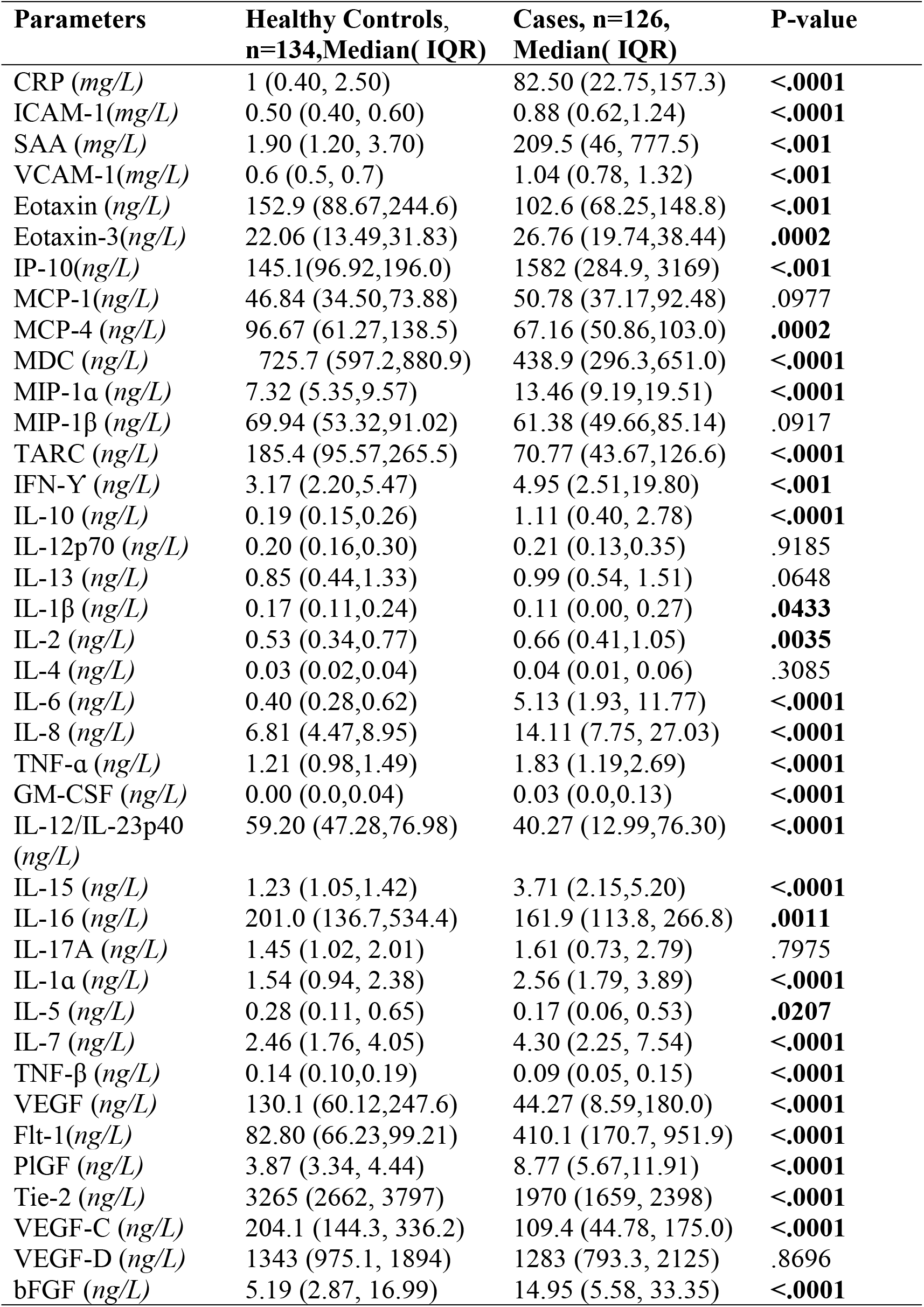
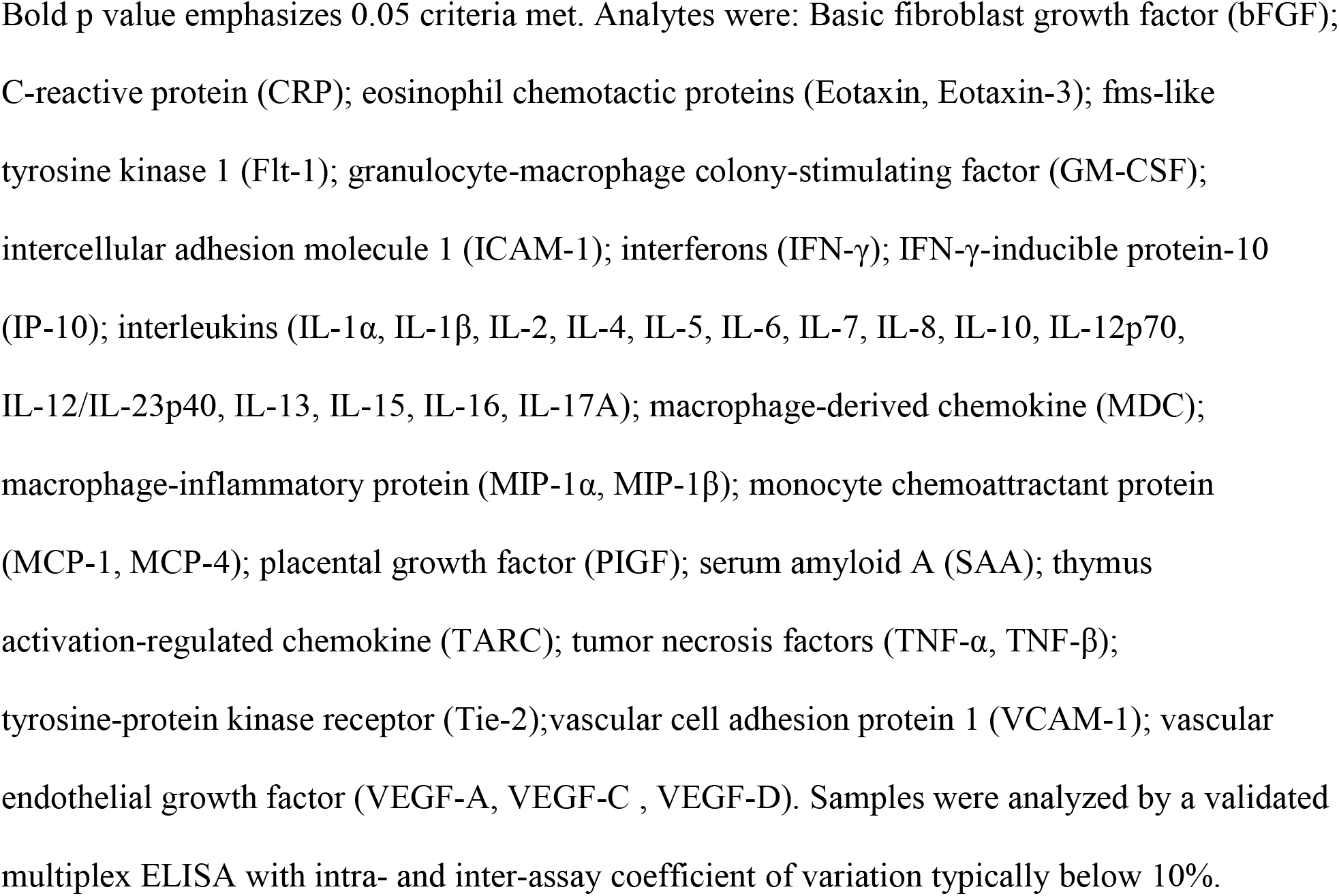
Concentration of inflammatory biomarkers sorted as healthy controls and COVID-19 patients

| Parameters | Healthy Controls,<br>n=134,Median( IQR) | Cases, n=126,<br>Median( IQR) | P-value |
| --- | --- | --- | --- |
| CRP (mg/L) | 1 (0.40, 2.50) | 82.50 (22.75,157.3) | <.0001 |
| ICAM-1(mg/L) | 0.50 (0.40, 0.60) | 0.88 (0.62,1.24) | <.0001 |
| SAA (mg/L) | 1.90 (1.20, 3.70) | 209.5 (46, 777.5) | <.001 |
| VCAM-1(mg/L) | 0.6 (0.5, 0.7) | 1.04 (0.78, 1.32) | <.001 |
| Eotaxin (ng/L) | 152.9 (88.67,244.6) | 102.6 (68.25,148.8) | <.001 |
| Eotaxin-3(ng/L) | 22.06 (13.49,31.83) | 26.76 (19.74,38.44) | .0002 |
| IP-10(ng/L) | 145.1(96.92,196.0) | 1582 (284.9, 3169) | <.001 |
| MCP-1(ng/L) | 46.84 (34.50,73.88) | 50.78 (37.17,92.48) | .0977 |
| MCP-4 (ng/L) | 96.67 (61.27,138.5) | 67.16 (50.86,103.0) | .0002 |
| MDC (ng/L) | 725.7 (597.2,880.9) | 438.9 (296.3,651.0) | <.0001 |
| MIP-1α (ng/L) | 7.32 (5.35,9.57) | 13.46 (9.19,19.51) | <.0001 |
| MIP-1β (ng/L) | 69.94 (53.32,91.02) | 61.38 (49.66,85.14) | .0917 |
| TARC (ng/L) | 185.4 (95.57,265.5) | 70.77 (43.67,126.6) | <.0001 |
| IFN-γ (ng/L) | 3.17 (2.20,5.47) | 4.95 (2.51,19.80) | <.001 |
| IL-10 (ng/L) | 0.19 (0.15,0.26) | 1.11 (0.40, 2.78) | <.0001 |
| IL-12p70 (ng/L) | 0.20 (0.16,0.30) | 0.21 (0.13,0.35) | .9185 |
| IL-13 (ng/L) | 0.85 (0.44,1.33) | 0.99 (0.54, 1.51) | .0648 |
| IL-1β (ng/L) | 0.17 (0.11,0.24) | 0.11 (0.00, 0.27) | .0433 |
| IL-2 (ng/L) | 0.53 (0.34,0.77) | 0.66 (0.41,1.05) | .0035 |
| IL-4 (ng/L) | 0.03 (0.02,0.04) | 0.04 (0.01, 0.06) | .3085 |
| IL-6 (ng/L) | 0.40 (0.28,0.62) | 5.13 (1.93, 11.77) | <.0001 |
| IL-8 (ng/L) | 6.81 (4.47,8.95) | 14.11 (7.75, 27.03) | <.0001 |
| TNF-α (ng/L) | 1.21 (0.98,1.49) | 1.83 (1.19,2.69) | <.0001 |
| GM-CSF (ng/L) | 0.00 (0.0,0.04) | 0.03 (0.0,0.13) | <.0001 |
| IL-12/IL-23p40 (ng/L) | 59.20 (47.28,76.98) | 40.27 (12.99,76.30) | <.0001 |
| IL-15 (ng/L) | 1.23 (1.05,1.42) | 3.71 (2.15,5.20) | <.0001 |
| IL-16 (ng/L) | 201.0 (136.7,534.4) | 161.9 (113.8, 266.8) | .0011 |
| IL-17A (ng/L) | 1.45 (1.02, 2.01) | 1.61 (0.73, 2.79) | .7975 |
| IL-1α (ng/L) | 1.54 (0.94, 2.38) | 2.56 (1.79, 3.89) | <.0001 |
| IL-5 (ng/L) | 0.28 (0.11, 0.65) | 0.17 (0.06, 0.53) | .0207 |
| IL-7 (ng/L) | 2.46 (1.76, 4.05) | 4.30 (2.25, 7.54) | <.0001 |
| TNF-β (ng/L) | 0.14 (0.10,0.19) | 0.09 (0.05, 0.15) | <.0001 |
| VEGF (ng/L) | 130.1 (60.12,247.6) | 44.27 (8.59,180.0) | <.0001 |
| Flt-1(ng/L) | 82.80 (66.23,99.21) | 410.1 (170.7, 951.9) | <.0001 |
| PlGF (ng/L) | 3.87 (3.34, 4.44) | 8.77 (5.67,11.91) | <.0001 |
| Tie-2 (ng/L) | 3265 (2662, 3797) | 1970 (1659, 2398) | <.0001 |
| VEGF-C (ng/L) | 204.1 (144.3, 336.2) | 109.4 (44.78, 175.0) | <.0001 |
| VEGF-D (ng/L) | 1343 (975.1, 1894) | 1283 (793.3, 2125) | .8696 |
| bFGF (ng/L) | 5.19 (2.87, 16.99) | 14.95 (5.58, 33.35) | <.0001 |

Bold p value emphasizes 0.05 criteria met. Analytes were: Basic fibroblast growth factor (bFGF); C-reactive protein (CRP); eosinophil chemotactic proteins (Eotaxin, Eotaxin-3); fms-like tyrosine kinase 1 (Flt-1); granulocyte-macrophage colony-stimulating factor (GM-CSF); intercellular adhesion molecule 1 (ICAM-1); interferons (IFN-γ ); IFN-γ-inducible protein-10 (IP-10); interleukins (IL-1α , IL-1β , IL-2, IL-4, IL-5, IL-6, IL-7, IL-8, IL-10, IL-12p70, IL-12/IL-23p40, IL-13, IL-15, IL-16, IL-17A); macrophage-derived chemokine (MDC); macrophage-inflammatory protein (MIP-1α , MIP-1β ); monocyte chemoattractant protein (MCP-1, MCP-4); placental growth factor (PIGF); serum amyloid A (SAA); thymus activation-regulated chemokine (TARC); tumor necrosis factors (TNF-α , TNF-β ); tyrosine-protein kinase receptor (Tie-2);vascular cell adhesion protein 1 (VCAM-1); vascular endothelial growth factor (VEGF-A, VEGF-C , VEGF-D). Samples were analyzed by a validated multiplex ELISA with intra- and inter-assay coefficient of variation typically below 10%.

Severe COVID-19 patients had significantly higher levels of CRP, ICAM-1, SAA, VCAM-1, IP-10, IL-10, IL-15, IL-16, IL-7, Flt-1 and VEGF-D than mild or moderate COVID-19 patients. MDC, IL-12/IL-23p40, IL-17A and TNF-β levels, on the other hand, were greater in mild or moderate COVID-19 patients than in severe COVID-19 patients (P < 0.05, Table 4). Between COVID-19 patients and healthy controls, median chemokine concentrations (MCP-1 and MIP-1β) were not statistically significant (P > 0.05, Table 3).

**TABLE 4.** Concentration of inflammatory biomarkers in mild/moderate versus severe COVID-19 patients

| Parameters | Mild/Moderate, n=58,<br>Median( IQR) | Severe, n=68,<br>Median( IQR) | P-value |
| --- | --- | --- | --- |
| CRP(mg/L) | **44.50 (9.0, 110.3)** | **99.50 (43.25, 187.0)** | **.0007** |
| ICAM-1(mg/L) | **0.74 (0.52, 1.14)** | **0.97 (0.67, 1.29 )** | **.0082** |
| SAA(mg/L) | **111.0 (19.25,335.5)** | **422.0 (139.0,883.3)** | **<.0001** |
| VCAM-1(mg/L) | **0.94 (0.72,1.22)** | **1.12 (0.86,1.37)** | **.0095** |
| Eotaxin (ng/L) | 114.7 (70.82,162.1) | 93.41(67.61,133.5) | .1945 |
| Eotaxin-3(ng/L) | 28.49 (19.03,37.54) | 25.47 (20.41,39.60) | .9388 |
| IP-10 (ng/L) | **471.5 (168.8,1826)** | **2084 (721.1,4228 )** | **<.0001** |
| MCP-1 (ng/L) | 55.51 (41.01 ,97.44 ) | 45.63 (35.55,87.95) | .1499 |
| MCP-4 (ng/L) | 60.40 (48.32 ,90.30 ) | 75.36 (53.26,111.6 ) | .1427 |
| MDC (ng/L) | **563.0 (326.1 ,762.9 )** | **380.4 (280.7,504.3 )** | **.0036** |
| MIP-1α (ng/L) | 13.06 (9.36 , 20.52 ) | 13.54 (8.57 ,19.39) | .8348 |
| MIP-1β (ng/L) | 61.01 (49.18, 81.78 ) | 61.70 (50.14,86.63 ) | .9204 |
| TARC (ng/L) | 68.64 (45.04, 180.0) | 71.94 (43.08,109.2 ) | .3889 |
| IFN-Y (ng/L) | 5.99 (2.98, 23.11) | 4.24 ( 1.87, 15.77) | .2639 |
| IL-10 (ng/L) | **0.60 (0.31, 2.18)** | **1.49 (0.55, 3.78)** | **.0025** |
| IL-12p70 (ng/L) | 0.21 (0.15, 0.34) | 0.24 (0.12, 0.38) | .7706 |
| IL-13 (ng/L) | 1.14 (0.76, 1.70) | 0.89 (0.50,1.43) | .0698 |
| IL-1β (ng/L) | 0.14 (0.0, 0.27) | 0.060 (0.0, 0.25) | .6639 |
| IL-2 (ng/L) | 0.71 (0.36, 1.03) | 0.61 (0.46, 1.08) | .4399 |
| IL-4 (ng/L) | 0.04 (0.02, 0.06) | 0.03 (0.01,0.06 ) | .7863 |
| IL-6 (ng/L) | 4.78 (1.58, 11.21) | 5.61 (2.55, 14.40) | .1715 |
| IL-8 (ng/L) | 13.83 (7.43, 26.27) | 14.23 (8.24, 27.71) | .9504 |
| TNF-α (ng/L) | 1.71 (1.17, 2.81) | 1.88 (1.22, 2.71) | .6076 |
| GM-CSF (ng/L) | 0.04 (0.0, 0.14) | 0.01 (0.0, 0.10) | .4333 |
| IL-12/IL-23p40 (ng/L) | **60.56 (28.58, 111.8)** | **29.90 (11.20, 52.28)** | **.0005** |
| IL-15 (ng/L) | **2.66 (1.68, 5.01)** | **4.03 (3.09, 5.22)** | **.0037** |
| IL-16 (ng/L) | **138.5 (98.65, 247.8)** | **179.1 (129.1, 271.5)** | **.0098** |
| IL-17A (ng/L) | **2.34 (1.15, 3.47)** | **0.94 (0.50,2.18)** | **.0001** |
| IL-1α (ng/L) | 2.43 (1.88, 4.07) | 2.68 (1.59, 3.87) | .9718 |
| IL-5 (ng/L) | **0.33 (0.16 , 0.76)** | **0.09 (0.0, 0.28)** | **<.0001** |
| IL-7 (ng/L) | **2.71 (1.78 , 5.27 )** | **5.69 (3.49 , 10.44)** | **<.0001** |
| TNF-β (ng/L) | **0.13 (0.09 ,0.17)** | **0.06 (0.04, 0.13 )** | **<.0001** |
| VEGF (ng/L) | 41.65 (7.83, 136.8) | 48.95 (8.89 , 191.6) | .5632 |
| Flt-1 (ng/L) | **201.6 (107.8, 423.2)** | **701.6 (331.2, 1783)** | **<.0001** |
| PlGF (ng/L) | 8.62 (5.10, 10.97) | 8.88 (6.25, 13.59) | .1669 |
| Tie-2 (ng/L) | 2091 (1680,2499) | 1923 (1644, 2229) | .3442 |
| VEGF-C (ng/L) | 109.4 (43.50,191.0) | 107.8 (46.16,169.2) | .8026 |
| VEGF-D (ng/L) | **1061 (746.3 ,1563)** | **1686 (922.4,2384)** | **.0071** |
| bFGF (ng/L) | 14.63 (4.43 ,34.33) | 15.31 (8.44 ,32.84) | .5251 |

Bold p value emphasizes 0.05 criteria met.

**TABLE 5.** Concentrations of inflammatory biomarkers in healthy controls and cases groups for adjusted age (≤ 40 years).

| Parameters | Healthy Controls,<br>n=126, Median ( IQR) | Cases, n=54,<br>Median( IQR) | P-value |
| --- | --- | --- | --- |
| CRP(mg/L) | 1(0.4, 2.4) | 67.50 (12.5,115.3) | **<.0001** |
| ICAM-1(mg/L) | 0.5 (0.4, 0.6) | 0.86 (0.5, 1.29 ) | **<.0001** |
| SAA(mg/L) | 1. 9 (1.2 , 3.6 ) | 131 (17 , 508.8 ) | **<.0001** |
| VCAM-1(mg/L) | 0.6 (0.5 , 0.7) | 0.9 (0.7 , 1.1) | **<.0001** |
| Eotaxin (ng/L) | 153.8 (89.3 ,244.6 ) | 90.7 (57.8 ,145.8 ) | **<.0001** |
| Eotaxin-3(ng/L) | 21.6 (13.4 , 32.2 ) | 26.8 (20.8 , 37.5) | **.0053** |
| IP-10 (ng/L) | 145.1 (96.9 , 191.8 ) | 463.4 (159.8,1944) | **<.0001** |
| MCP-1 (ng/L) | 45.9 ( 34.5 , 72.4 ) | 45.6 (35.6 , 86.5 ) | .6157 |
| MCP-4 (ng/L) | 92.2 (59.8 , 138.2 ) | 60.6 (49.8 , 92.8 ) | **.0005** |
| MDC (ng/L) | 723.7 (600.9 ,876.1 ) | 501.1 (308.7,743) | **<.0001** |
| MIP-1α (ng/L) | 7.4 (5.3 , 9.6 ) | 12.9 (9.1 , 18.3) | **<.0001** |
| MIP-1β (ng/L) | 69.9 (53.9 , 87.6 ) | 59.6 (48 , 81.8) | **.0460** |
| TARC (ng/L) | 187.7 (106.5 , 265.5 ) | 63.7 (41.6 ,152.9 ) | **<.0001** |
| IFN-Y (ng/L) | 3.2 (2.2 , 5.5 ) | 4 ( 1.7 , 19.1) | **.0376** |
| IL-10 (ng/L) | 0.2 (0.1, 0.3) | 0.8 (0.3, 2.7) | **<.0001** |
| IL-12p70 (ng/L) | 0.2 (0.2, 0.3) | 0.2 (0.2, 0.4) | .3699 |
| IL-13 (ng/L) | 0.8 (0.4, 1.3) | 1.1 (0.8, 1.7) | **.0036** |
| IL-1β (ng/L) | 0.2 (0.1, 0.2) | 0.1 (0.0, 0.3) | .0974 |
| IL-2 (ng/L) | 0.5 (0.3, 0.8) | 0.7 (0.5, 1.1) | **.0016** |
| IL-4 (ng/L) | 0.03 (0.02, 0.04) | 0.04 (0.02, 0.06) | .1804 |
| IL-6 (ng/L) | 0.4 (0.3, 0.6) | 3.6 (1.8, 10.7) | **<.0001** |
| IL-8 (ng/L) | 6.8 (4.4, 9.3) | 11.1 (6.1, 34.2) | **<.0001** |
| TNF-α (ng/L) | 1.2 (0.9, 1.5) | 1.6 (1.1, 2.6) | **.0012** |
| GM-CSF (ng/L) | 0.0 (0.0, 0.04) | 0.04 (0.0, 0.2) | **.0002** |
| IL-12/IL-23p40 (ng/L) | 59.2 (47.3, 77.2) | 54.4 (13.4, 84.8) | .1528 |
| IL-15 (ng/L) | 1.2 (1, 1.4) | 2.9 (1.8, 5.2) | **<.0001** |
| IL-16 (ng/L) | 200 (134.3, 590.3) | 126.5 (94.3, 202.7) | **<.0001** |
| IL-17A (ng/L) | 1.5 (1, 2) | 1.9 (0.9, 3.4) | .2010 |
| IL-1α (ng/L) | 1.5 (0.9, 2.4) | 2.5 (1.9, 3.5) | **<.0001** |
| IL-5 (ng/L) | 0.28 (0.1, 0.6) | 0.26 (0.1, 0.7) | .7479 |
| IL-7 (ng/L) | 2.5 (1.8, 4.1) | 3.3 (1.9, 5.8) | .0595 |
| TNF-β (ng/L) | 0.14 (0.1, 0.2) | 0.12 (0.07, 0.16) | **.0283** |
| VEGF (ng/L) | 127.6 (60.1, 238.1) | 41.7 (12.4, 167.8) | **<.0001** |
| Flt-1 (ng/L) | 83.2 (66.1, 100.7) | 214.2 (136.5, 618.) | **<.0001** |
| PlGF (ng/L) | 3.9 (3.3, 4.4) | 7.8 (4.9, 11.2) | **<.0001** |
| Tie-2 (ng/L) | 3284 (2683, 3845) | 2098 (1731, 2549) | **<.0001** |
| VEGF-C (ng/L) | 206.5 (146.3, 324.3) | 113.5 (45.1, 195.2) | **<.0001** |
| VEGF-D (ng/L) | 1335 (975.1, 1882) | 1493 (918.4, 2496) | .1389 |
| bFGF (ng/L) | 5.3 (2.9, 17.1) | 12.6 (4.1, 31.3) | **.0208** |

Bold p value emphasizes 0.05 criteria met.

Based on the correlations of 30 significantly elevated cytokines and chemokines; CRP, SAA, IL-6, IL-8, IP-10, TNF-α, IL-10, VEGF-D, MDC and VCAM-1 were the ones that discriminated between mild or moderate and severe cases (Table 6). The diagnostic value of cytokines and chemokines for illness severity was assessed using the receiver operating characteristic (ROC) curve and the area under the ROC curve (AUC) (Table 7 and Figure 1). AUC (95 percent confidence intervals) were CRP 0.9817 (0.9703-0.9931), SAA 0.9783 (0.9610-0.9956), IL-6 0.9634 (0.9425-0.9843), IL-8 0.7800 (0.7238-0.8362), IP-10 0.8735 (0.8265-0.9204), TNF-α 0.7168 (0.6521-0.7816), IL-10 0.8793 (0.8328-0.9257), VEGF-D 0.5059 (0.4347-0.5772), MDC 0.7894 (0.7324-0.8463) and VCAM-1 0.8465 (0.7960- 0.8971) respectively (P < 0.05). SAA, CRP and IL-6 have very high sensitivity and specificity (Figure 1 and Table 7). TNF-α and VEGF-D have poor sensitivity and specificity when compared to CRP, SAA and IL-6 (Figure 1). The hierarchy among the 10 selected cytokines and chemokines in distinguishing between healthy controls and COVID-19 patients, as well as between mild or moderate and severe COVID-19 patients, was validated by ROC and AUC analyses (Table 7 & 8). Figure 2 demonstrates the use of a ROC curve to distinguish between mild/moderate and severe COVID-19 cases for severity prediction.

**FIG 1.**
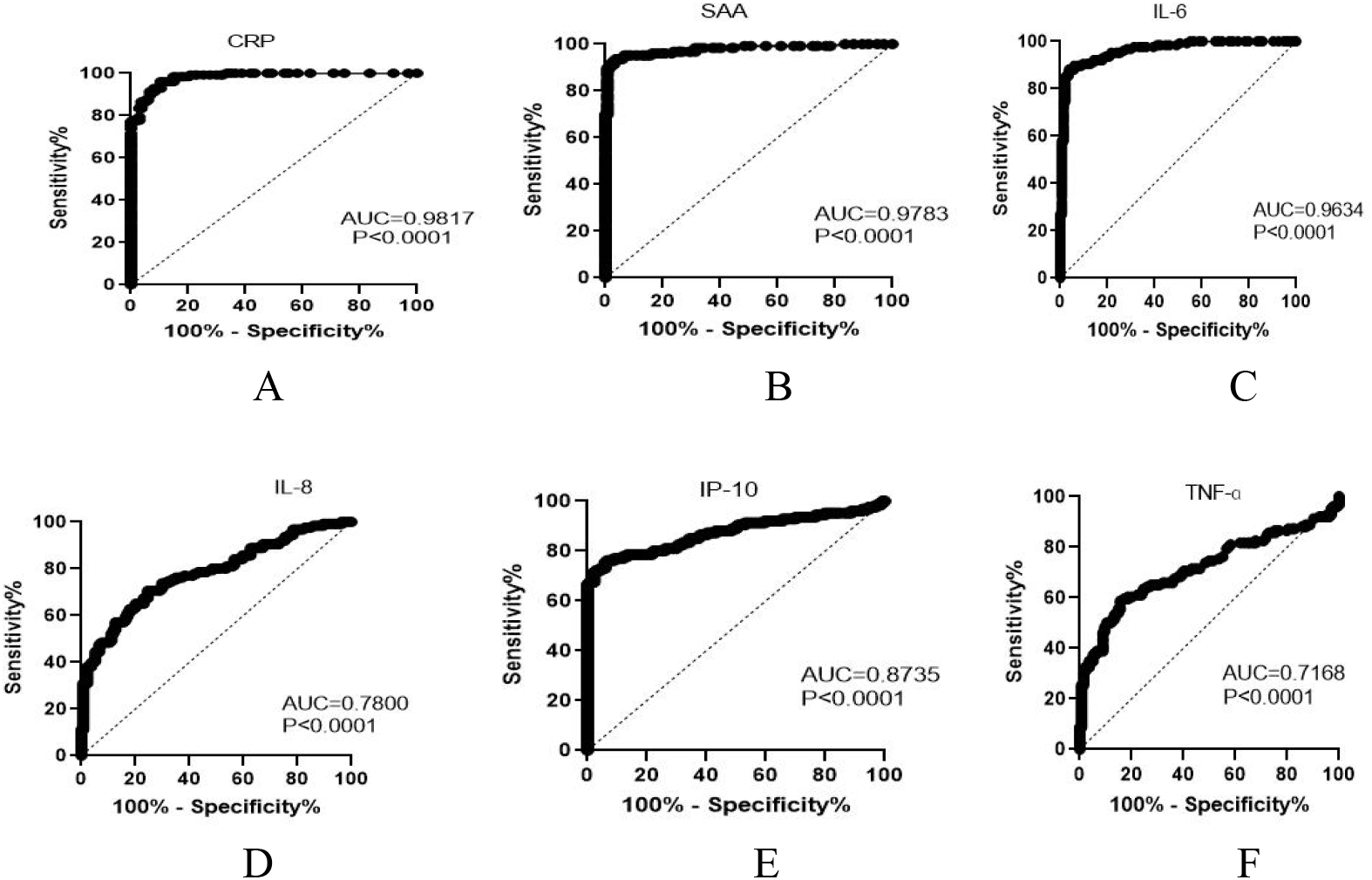
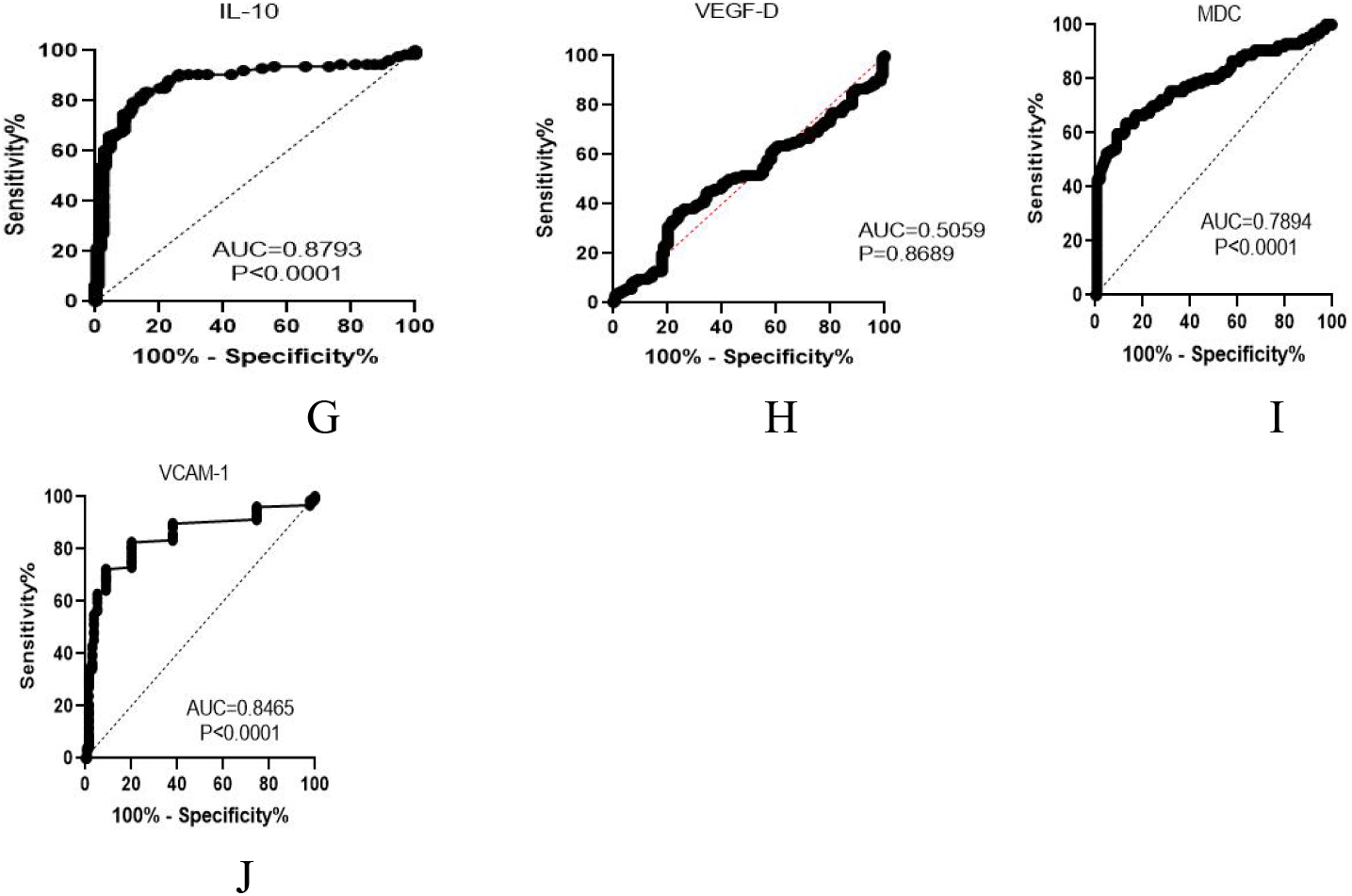
Receiver operating characteristic analysis of healthy control subjects versus COVID-19 patients, using specific biomarkers A) CRP, C-reactive protein. B) SAA, serum amyloid A. C) IL-6, interleukin 6. D) IL-8, interleukin 8. E) IP-10, interferon-γ-inducible protein-10. F) TNF-α, tumor necrosis-α. G) IL-10, interleukin-10. H) VEGF-D, vascular endothelial growth factor-D. I) MDC, macrophage-derived chemokine (MDC). J) VCAM-1, vascular cell adhesion protein 1.

**FIG 2.**
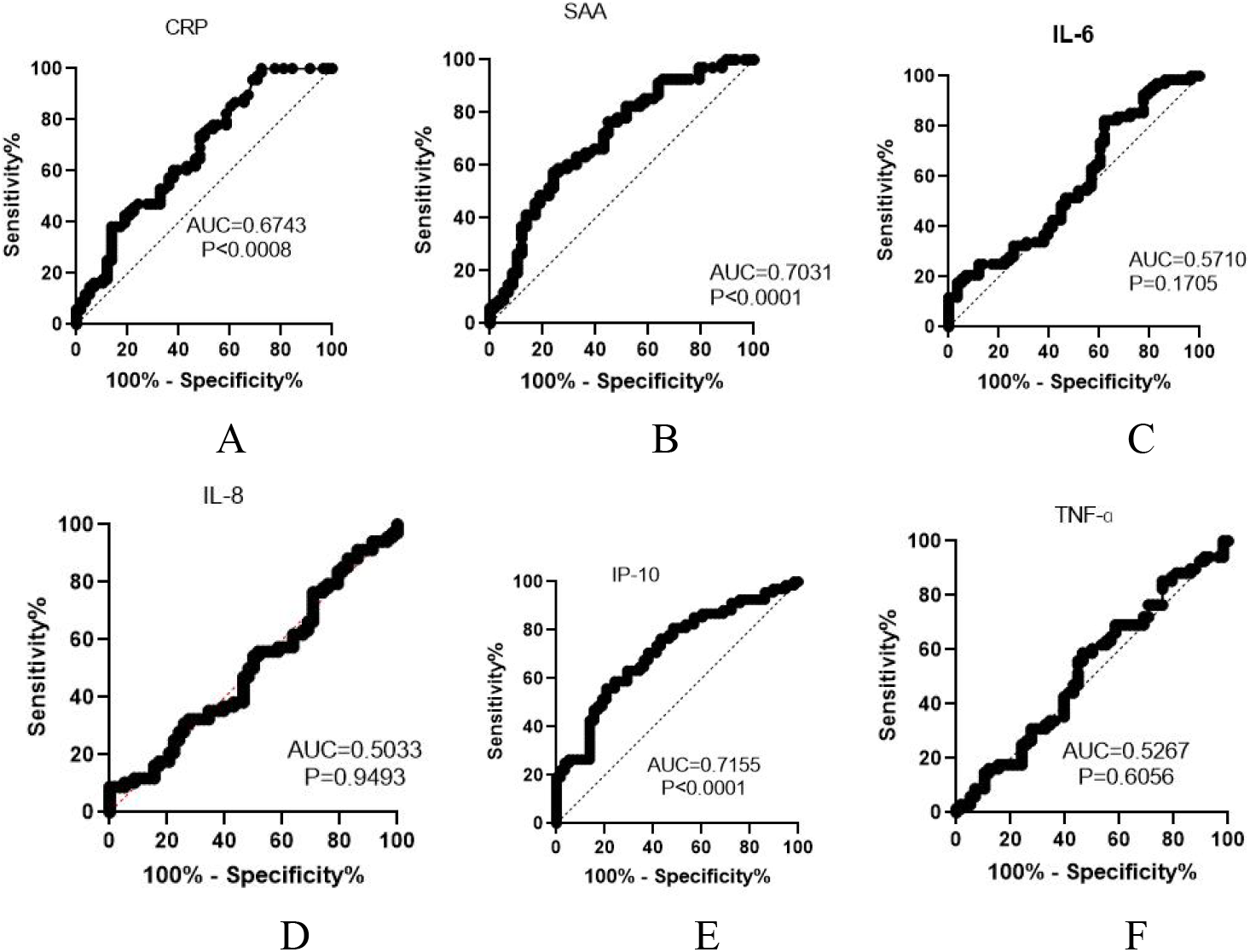
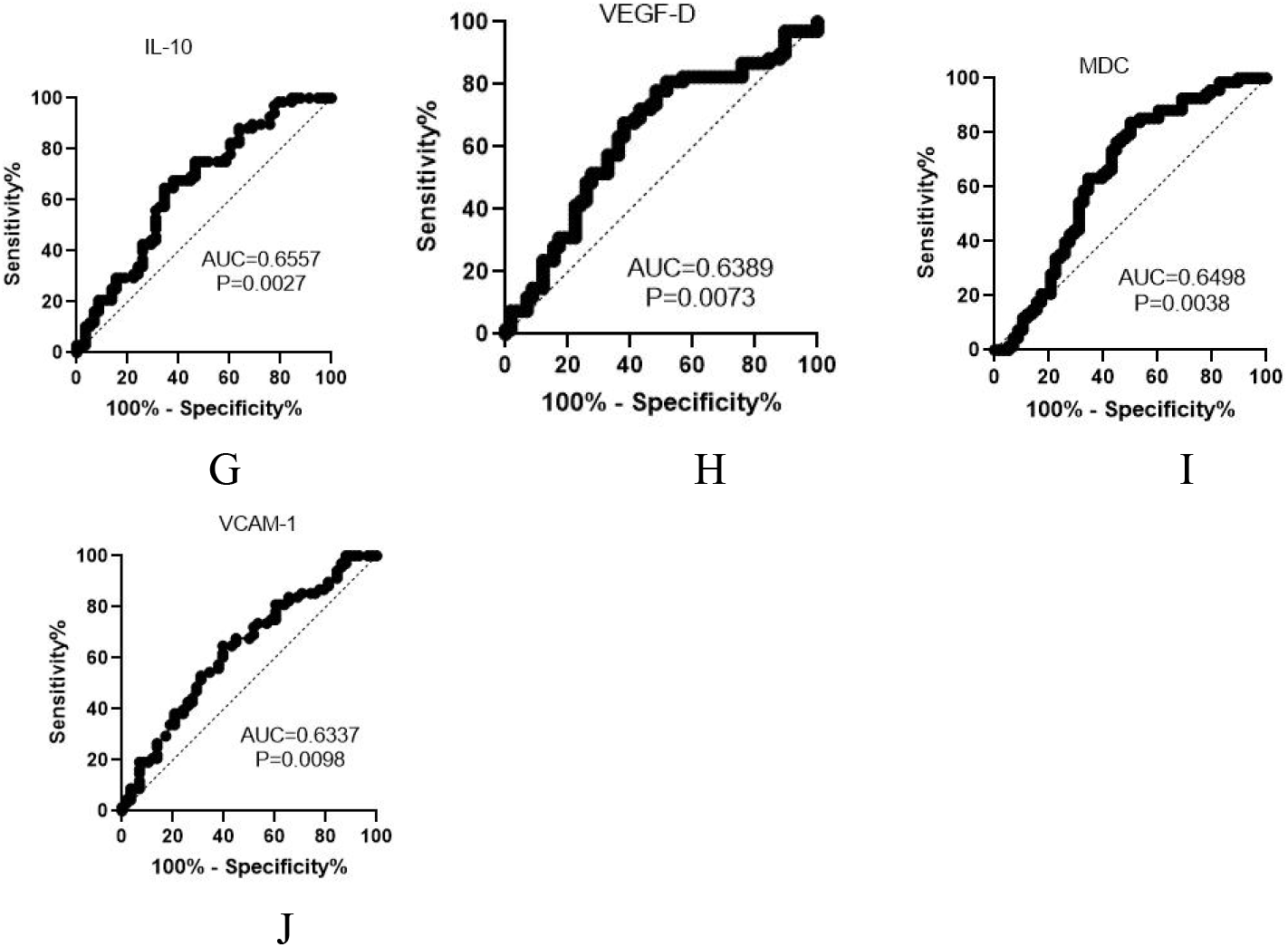
Receiver operating characteristic curve of indicators between mild/moderate and severe COVID-19 for the prediction of severity. A) CRP, C-reactive protein. B) SAA, serum amyloid A. C) IL-6, interleukin 6. D) IL-8, interleukin 8. E) IP-10, interferon-γ-inducible protein-10. F) TNF-α, tumor necrosis-α. G) IL-10, interleukin-10. H) VEGF-D, vascular endothelial growth factor-D. I) MDC, macrophage-derived chemokine (MDC). J) VCAM-1, vascular cell adhesion protein 1.

**TABLE 6.**
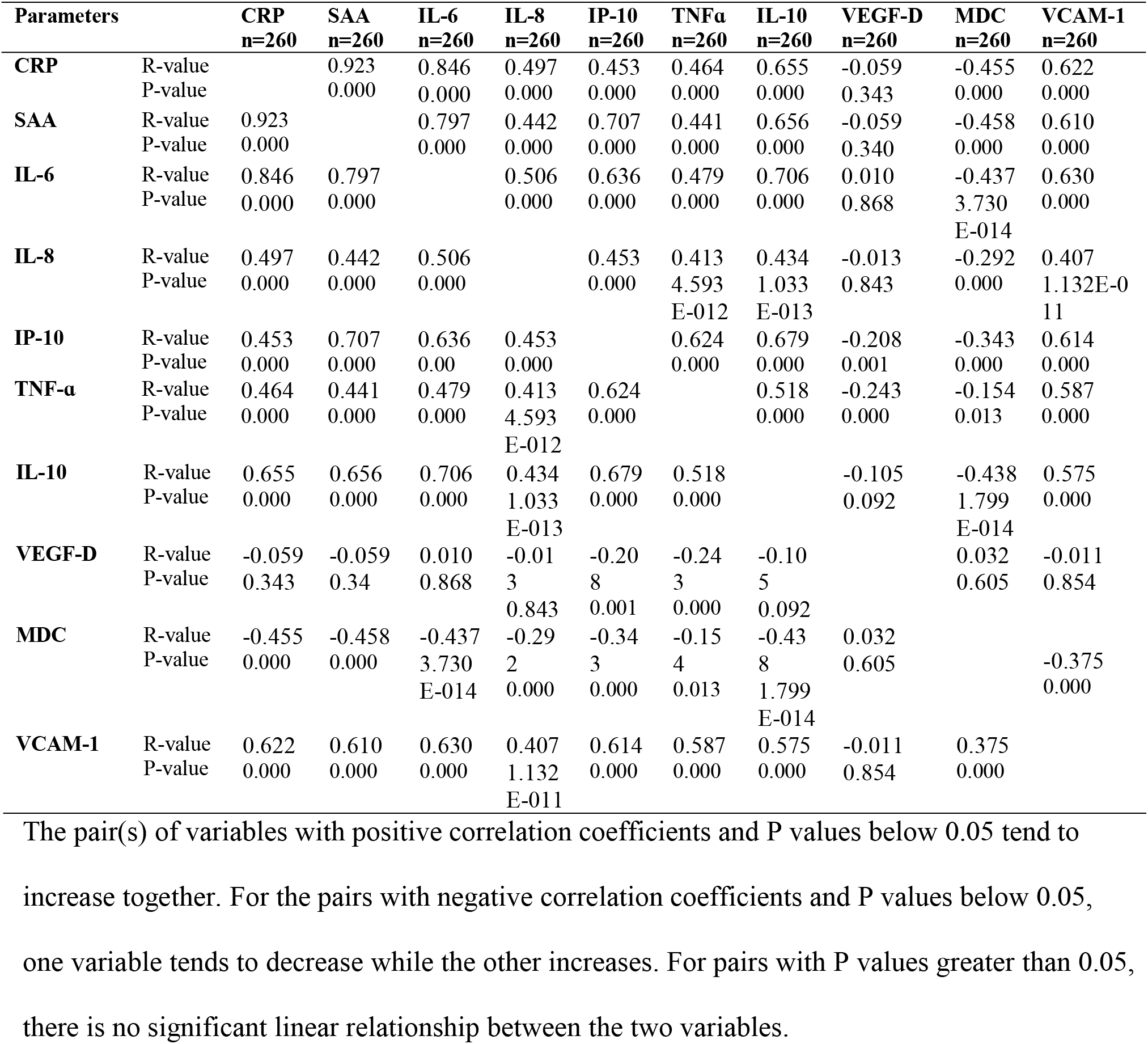
Correlations of cytokines and chemokines. Data from all subjects (healthy controls, mild/moderate and severe cases).

**TABLE 7.** Area under the receiver operating characteristic curve, sensitivity, specificity & cutoff values of specific inflammatory markers of distinguishing healthy controls and cases

| Variable | AUC (95 % CI) | Sensitivity %<br>(95 % CI) | Specificity%<br>(95 % CI) | Cutoff | P-value |
| --- | --- | --- | --- | --- | --- |
| CRP | 0.9817(0.9703-0.9931) | 75.4(67.2-82.1) | 100(97.2- 100) | > 22.50 | **<.0001** |
| SAA | 0.9783(0.9610-0.9956) | 88.9(82.2- 93.3) | 99.3(95.9-99.9) | > 11.50 | **<.0001** |
| IL-6 | 0.9634(0.9425-0.9843) | 82.5(74.9- 88.2) | 97.8(93.6- 99.4) | > 1.49 | **<.0001** |
| IL-8 | 0.7800(0.7238-0.8362) | 38.1(30.1- 46.8) | 97.8(93.6- 99.4) | > 18.92 | **<.0001** |
| IP-10 | 0.8735(0.8265-0.9204) | 69.8(61.3- 77.2) | 97.8(93.6- 99.4) | > 374.2 | **<.0001** |
| TNF-α | 0.7168(0.6521-0.7816) | 30.9(23.5- 39.5) | 97.8(93.6- 99.4) | > 2.505 | **<.0001** |
| IL-10 | 0.8793(0.8328- 0.9257) | 53.2(44.5- 61.7) | 97.8(93.6- 99.4) | > 1.010 | **<.0001** |
| VEGF-D | 0.5059(0.4347- 0.5772) | 3.9(1.7- 8.9) | 97.8(93.6- 99.4) | < 429.9 | .8689 |
| MDC | 0.7894(0.7324- 0.8463) | 46.8(38.3- 55.5) | 97(92.6 - 98.8) | < 418.8 | **<.0001** |
| VCAM-1 | 0.8465(0.7960- 0.8971) | 32.5(24.9- 41.1) | 98.5(94.7- 99.7) | > 1.215 | **<.0001** |

Bold p value emphasizes 0.05 criteria met. Cut off value units for CRP, SAA and VCAM-1 were (mg/L) and IL-6, IL-8, IP-10, TNF-α, IL-10, VEGF-D and MDC were (ng/L).

**TABLE 8.** Area under the receiver operating characteristic curve, sensitivity, specificity & cutoff values of specific markers in discriminating mild/moderate and severe COVID-19 patients

| Variable | AUC (95 % CI) | Sensitivity %<br>(95 % CI) | Specificity%<br>(95 % CI) | Cutoff | P-value |
| --- | --- | --- | --- | --- | --- |
| CRP | 0.6743(0.5799- 0.7687) | 7.4(3.2 - 16.1) | 98.3(90.9-99.9) | > 395 | **0.0008** |
| SAA | 0.7031(0.6115 -0.7947) | 5.9(2.3- 14.2) | 98.3(90.9- 99.9) | > 1074 | **<0.0001** |
| IL-6 | 0.5710(0.4696 -0.6724) | 11.8(6.1- 21.5) | 100(93.8- 100) | > 77.72 | 0.1705 |
| IL-8 | 0.5033(0.4015- 0.6051) | 7.4(3.2- 16.1) | 100(93.8 - 100) | > 305.7 | 0.9493 |
| IP-10 | 0.7155(0.6263- 0.8047) | 20.6(12.7- 31.6) | 98.3(90.9- 99.9) | > 5185 | **<0.0001** |
| TNF-α | 0.5267(0.4246 -0.6289) | 1.5(0.1- 7.9) | 98.3(90.9- 99.9) | > 36.32 | 0.6056 |
| IL-10 | 0.6557(0.5589- 0.7524) | 2.9(0.5 - 10.1) | 100(93.8- 100) | > 46.46 | **0.0027** |
| VEGF-D | 0.6389(0.5404-0.7375) | 7.4(3.2 -16.1) | 96.6(88.3- 99.4) | < 482.3 | **0.0073** |
| MDC | 0.6498(0.5495-0.7502) | 4.4(1.2- 12.2) | 93.1(83.6- 97.3) | < 156.1 | **0.0038** |
| VCAM-1 | 0.6337(0.5364 -0.7311) | 2.9(0.5- 10.1) | 98.3(90.9- 99.9) | > 2.120 | **0.0098** |

Bold p value emphasizes 0.05 criteria met. Cut off value units for CRP, SAA and VCAM-1 were (mg/L) and IL-6, IL-8, IP-10, TNF-α , IL-10, VEGF-D and MDC were (ng/L).

Table 9 shows a binary logistic regression analysis that links CRP and age to COVID-19 disease severity. These findings indicated that CRP and age were statistically significant (P < 0.05).

**TABLE 9.** Binary logistic regression analysis indicating factors associated with disease severity

| Variable | AOR(95% CI) | P-value |
| --- | --- | --- |
| Age | 1.056 (1.033-1.081) | <0.001 |
| CRP | 1.004(1.000-1.008) | 0.044 |

AOR=Adjusted odds ratio; co-morbidity and gender were insignificant with p-value ( 0.696 and 0.730 ).

## DISCUSSION

This study looks at 39 cytokines and chemokines in COVID-19 patients’ inflammation and disease severity. SARS-COV-2 infection can cause varied inflammatory response abnormalities. The pathogenesis of many viral infections, including coronaviruses, is connected to aberrant virus-induced immune responses (15). Immune responses, particularly expression of cytokines and chemokines, are closely regulated in normal physiology; deviations from this control might result in different biochemical and clinical manifestations (16). This study is the first comparative study of COVID-19 patients in Ethiopia, to the best of our knowledge, to investigate different cytokine and chemokine profiles in discriminating between healthy controls and COVID-19 patients; mild/moderate and severe cases and prognosis, classification and predicting disease severity.

The average age of the severe group was much higher than mild/moderate group, in line with earlier research (17). This showed that age could be a risk factor for COVID-19 severity and it was assumed that ACE2 density was shown to be positively connected with age (18; 19). Gender distribution in the group was not statistically significant (Table 1). However, male patients were more likely to develop severe illness in the severe group than in the other groups. This could be due to gender linked immunological differences (20).

Pro-inflammatory cytokines and interferons (IFNs) were shown to be greater in COVID-19 patients (21). These cytokines aid in the elimination of infections as well as the maintenance of cellular homeostasis. Dysregulated pro-inflammatory cytokine release, on the other hand, contributes to cytokine storm, a potentially fatal condition caused by inflammatory cells’ excessive cytokine production (22). Inflammatory cytokine and chemokine levels were found to be significantly higher in patients than in the healthy controls group. This study consistent with a prior study comparing patients with severe COVID-19 to those with mild disease (23). In comparison to mild/moderate cases of COVID-19, plasma from patients with severe cases showed a greater tendency for levels of IL-6, IL-8 and TNF-α indicating a pro-inflammatory response. This research supports the findings of the prior research (24).

C-reactive protein (CRP) is increased in response to IL-6, IL-1β and TNF-α activation during infection and systemic inflammation (25). CRP levels were significantly greater in the severe group compared to the other groups in this investigation, which is consistent with a prior study that found CRP levels may represent disease severity in COVID-19 patients (26, 27). CRP is a non-specific measure used to distinguish between infectious and non-infectious diseases caused by viruses or bacteria (28, 29). Many studies suggest that CRP could be used to predict prognosis even before the onset of sickness as a biomarker for COVID-19 severity and that an increase in CRP is linked to a poor COVID-19 prognosis (27). It can also be used as an early indicator of infection and inflammation. According to the current study findings and previous researches, increased levels of CRP have clinical diagnostic and prognostic value during COVID-19 infection.

Endothelial cells aid in the recruitment of leukocytes from the circulation to infection and inflammatory sites (30). Inflammatory cytokines initiate different kinase cascades during SARS-CoV-2 infection, leading to the activation of transcription factors like NF-β, which in turn stimulate adhesion molecules like ICAM-1 and VCAM-1. ICAM-1 promotes leukocyte arrest and firm adhesion, as well as monocyte and lymphocyte transmigration. In the current study, COVID-19 patients had higher levels of ICAM-1 and VCAM-1 than healthy control and the severe group had higher levels than the mild or moderate group, which was consistent with earlier findings (31). This suggests that ICAM-1 and VCAM-1 could be used as a biomarker to predict the severity of COVID-19 disease and may have a role in coagulation dysfunction. In line with the previous study, SAA levels in the current study were considerably higher in the COVID-19 patients than healthy controls. SAA is released by hepatocyte cells in response to inflammatory cytokines such as IL-6, TNF-α and IL-1β during an acute phase response. One of the most common clinical symptoms of severe COVID-19 is ARDS (32). There were patients with ARDS who had a considerably higher level of SAA. SAA could be used as a biomarker to track the evolution of respiratory disorders like COVID-19, according to our findings. The median levels of both CRP and SAA were greater in COVID-19 patients than healthy controls after age adjustment (comparing age ≤ 40 years in both COVID-19 patients and healthy controls). CRP and SAA were both linked to disease severity; have a strong and positive linear correlation. Since measurement of different cytokines/chemokines is time-consuming and costly, correlation analysis aids clinical prediction of one biomarker in terms of another. Thus, correlations were performed in the current investigation, implying that plasma levels of CRP, SAA, IL-6, IP-10, VCAM-1, and IL-10, among the linked, exhibit a significant and beneficial association with COVID-19 patient progression. CRP, SAA, IL-6, IP-10, VCAM-1, TNF-α and IL-10 all had a negative relationship with plasma levels of VEGF-D and MDC.

In this study, in individuals with COVID-19, IP-10 levels were higher than in healthy controls and the same is true after age adjustment. Furthermore, when the viral infection was linked with a pulmonary pathology (e.g., influenza and SARS-COV-2), IP-10 levels were higher than when the infection was associated with a non-pulmonary pathology (e.g., human rhinovirus) (13). IP-10 levels were significantly greater in severe SARS-CoV-2-positive patients than in mild or moderate patients, implying that IP-10 could be a useful biomarker for distinguishing between mild/moderate and severe COVID-19 patients and predicting disease severity.

In SARS, Middle East respiratory syndrome-CoV and SARS-CoV-2 infected patients, excessive cytokine production has been associated to pulmonary inflammation and acute lung damage (15, 33). IL-10 is an anti-inflammatory cytokine that was shown to be higher in COVID-19 patients with severe illness in this study.This was in line with the findings of a previous study (23). It is important for immune system homeostasis. IL-10 is a cytokine with many functions that reduces the inflammatory response by direct effect on macrophages and T and B-cells (34). IL-10 is known to produce T-cell anergy or non-responsiveness in anti-tumour cell responses as well as viral infection (35). As a result, increased IL-10 levels in severe COVID-19 patients were initially attributed to a negative feedback mechanism which is suggestive of anti-inflammatory response (36). Furthermore, as compared to healthy controls and mild/moderate COVID-19 patients, plasma IL-7 levels were considerably higher in severe COVID-19 patients in this study.

This research supports the findings of the prior research (37). T-cell growth and functions are dependent on IL-7. Increased levels of IL-7 in the blood have been linked to a reduction in the number of T cells in the body (38). As a feedback response to lymphopenia, serum IL-7 concentration was also negatively related to the number of T cells, CD4^+^and CD8^+^ cells, highlighting the link between lymphopenia and higher IL-7 levels in SARS-CoV-2 infection (39).

In this study, mild/ moderate COVID-19 cases had considerably greater levels of IL-12/IL-23p40 than severe COVID-19 patients and healthy controls. This is in line with the findings of the prior study (40). Induction of IL-12 is essential to sustain NK cell counts during the early stages of SARS-CoV-2 infection and this induction could aid in evasion from virus spreading. The number of peripheral NK cells in patients with severe COVID-19 was much lower than in healthy people (41). Furthermore, IL-17A is a homeostatic proinflammatory cytokine that is also involved in the development of autoimmune disorders. Our findings revealed that IL-17A levels in severe COVID-19 patients were significantly lower than in mild or moderate COVID-19 patients. A meta-analysis comparing IL-17A levels in severe and moderate patients found that severe patients had greater levels (42). The discrepancy could be due to the data limitations that were revealed during the meta analysis.

Similarly, TGF-β levels were found to be decreased in severe patients compared to mild or moderate patients and healthy controls (Table 3 and 4). This is consistent with the concept that serum TGF-β levels peak during the first two weeks of acute COVID-19 infection and limit NK cell function in a TGF-dependent way (43). As a result, premature TGF-β production is thought to be a characteristic of severe COVID-19. TGF-β is released from a variety of sources in response to SARS-CoV-2 infection, including dysregulated coagulation and fibrinolytic pathways; neutrophils infiltrating the lungs in large numbers and macrophages migrating to the lungs to phagocytize apoptotic bronchial epithelial cells, pneumocytes, T-lymphocytes and neutrophils (44). The potential of this cytokine to recruit more neutrophils and remodel the airways through modulating processes used by the virus to generate pulmonary fibrosis explains its influence in SARS-CoV-2 infection.

The severe group had significantly greater levels of VEGF-D than the mild or moderate group. There was a data shortage for this biomarker, but it was consistent with the prior research, which compared critical and severe group (45). We hypothesized that a high level of VEGF-D is linked to a storm of blood clots in COVID-19 patients, which eventually leads to disease severity. The ROC curve data further revealed that CRP, SAA and IL-6 levels had excellent sensitivity and specificity for COVID-19 severity. This biomarker’s likely hood ratio value also accurately predicted illness dynamics, making it easier to recognize and act with underlying severe patients earlier, which was critical for lowering mortality. Binary logistic regression analysis found odds ratio (OR=1.056) for age, implying that the chance of being severe increases by 1.056 times with each unit rise in age. As people become older, their chance of becoming severely ill will increase. The same is true for CRP, with an OR of 1.004; one unit CRP increase has a 1.004 times increased likelihood of being severe. Increases in age and CRP will enhance the likelihood of severity, according to the regression model. These data suggest that both increment of CRP and age has a role in the advancement of COVID-19 disease severity.

The limitation of this study might be that it was impossible to get a comparable group because there were no elders in the healthy controls group (for age > 40 years), though adjustments were tried. Because blood was tapped and severity assessed shortly after positive test results, often at an outpatient setting, the study could not identify any future disease progression after the date of blood draw. Characterizing immune response is crucial for guiding public health interventions and measures. Thirty different cytokine and chemokine levels were shown to be elevated as a result of the severity of the disease in this investigation, suggesting that they could be utilized to predict the severity and prognosis of COVID-19 patients. Among elevated plasma cytokine/chemokine levels; CRP, ICAM-1, SAA, VCAM-1, IP-10, IL-7, IL-10 and Flt-1 may suggest the severity of COVID-19. Increment of these cytokines and chemokines in COVID-19 patients indicate an abnormal immune response and the emergence of a cytokine storm, which worsens the disease and may lead to death. As a result, the critical biomarkers determined in this study may aid in clinical care monitoring and COVID-19 treatment follow up.

## Materials and Methods

### Study design and setting

A comparative study was conducted at Tikur Anbessa Specialized Hospital (TASH), Addis Ababa University, Ethiopia from January 27, 2021 to December 30, 2021. The hospital has 700 beds and serves over 500,000 patients in the outpatient department and 370,000-400,000 inpatients every year. It’s also one of the COVID-19 diagnosis and treatment centers.

### Study participants

The study included 126 COVID-19 patients admitted to TASH with laboratory-confirmed and clinically diagnosed COVID-19 using a convenient sampling method and 134 healthy controls who had no contact with known COVID-19 suspected individuals for at least 14 days and had a negative RT-PCR. All participants were 18 years and older. Healthy control participants were selected from administrative staff and medical students at College of Health Sciences. Participants who were anemic, pregnant or had taken a corticosteroid or immunosuppressant within 14 days of admission were all excluded from the study.

### Specimen collection

Five millilitre of venous blood was tapped by syringe and needle and EDTA-plasma was separated from whole blood and immediately frozen at −80°c until assay.

### Cytokine and chemokine analysis

A human cytokine 39-plex ELISA assay was used (Meso Scale Diagnostics, Rockville, MD, USA) to measure basic fibroblast growth factor (bFGF); C-reactive protein (CRP); eosinophil chemotactic proteins (Eotaxin, Eotaxin-3); fms-like tyrosine kinase 1 (Flt-1); granulocyte-macrophage colony-stimulating factor (GM-CSF); intercellular adhesion molecule 1 (ICAM-1); interferons (IFN-γ); IFN-γ-inducible protein-10 (IP-10); interleukins (IL-1α, IL-1β, IL-2, IL-4, IL-5, IL-6, IL-7, IL-8, IL-10, IL-12p70, IL-12/IL-23p40, IL-13, IL-15, IL-16, IL-17A); macrophage-derived chemokine (MDC); macrophage-inflammatory protein (MIP-1α, MIP-1β); monocyte chemoattractant protein (MCP-1, MCP-4), placental growth factor (PIGF); serum amyloid A (SAA); thymus activation-regulated chemokine (TARC); tumor necrosis factors (TNF-α, TNF-β); tyrosine-protein kinase receptor (Tie-2); vascular cell adhesion protein 1 (VCAM-1); vascular endothelial growth factor (VEGF-A, VEGF-C, VEGF-D) at Department of Medical Sciences, Gastroenterology and Hepatology Unit, Uppsala University, Sweden (14).

### Statistical analysis

IBM SPSS version 25.0 (Chicago, IL) and Prism version 8 were used. Descriptive summary measures; frequency, percentages, mean (SD) and median (IQR) were used to describe participants. Chi-square test was used to determine associations between categorical variables. Shapiro-Wilk test was used to determine normality. Mann-Whitney U test was used to compare biomarker concentrations by group. Binary logistic regression was used to identify predictors of disease severity. Receiver operating characteristics (ROC) curve was employed to set cutoff points to predict disease status and disease severity. In addition Spearman rank correlation was done to see correlation between cytokines/chemokines. P-value < 0.05 considered as statistically significant.

### Ethical considerations

The study was approved by departmental research ethics and review committee of the Department of Biochemistry, CHS, AAU (Ref.No.SoM/BCHM/068/2013) and by the institutional review board of College of Health Sciences, Addis Ababa University (Meeting 01/2021, Protocol 004/21/Biochem). The study was also approved by the national research ethics review committee at the Ministry of Science and Higher Education, Ethiopia (Ref.No.MoSHE/04/246/837/21). Permission to conduct the study was also obtained from TASH. All participants in the study (both COVID-19 patients and healthy controls) signed a written informed consent form.

## ACKNOWLEDGEMENTS

A.T sincerely acknowledges support from CDT-Africa of Addis Ababa University and Professor Per M. Hellström and Dr. Dominic-Luc Webb, Department of Medical Sciences, Gastroenterology and Hepatology Unit, Uppsala University, Sweden DLW gratefully acknowledges support from Selander’s Foundation (2020 & 2021) as well as OE and Edla Johansson’s Science Foundation (2017 & 2018). DLW & PMH gratefully acknowledge funding from ALF-medel (Uppsala-Örebro region, ALF-899261) and Vetenskapsrådet (2017-02243).

## AUTHOR CONTRIBUTIONS

Credit roles were as follows:

A.T: Design; preparation; data curation; data analysis; investigation; Methodology; Roles/Writing - original draft; Writing - review, editing & correction.

S. G: Design; Conceptualization; Formal analysis; Investigation; Methodology; Project administration; Resources; Software; Visualization; Roles/Writing - review & editing.

T. H: clinical investigation, Methodology, Supervision; Writing - review & editing.

T.M: Funding acquisition, Methodology; Project administration; Resources; Supervision; Writing - review & editing.

T.G: Performed statistics and produced tables and figures.

PMH: Conceptualization; Funding acquisition; Investigation; Methodology; Project administration; Resources; Supervision; Writing - review & editing.

DLW: Conceptualization; Data curation; Formal analysis; Funding acquisition; Investigation; Methodology; Project administration; Resources; Software; Supervision; Validation; Visualization; Roles/Writing - original draft; Writing - review & editing.

A.T, S.G, T.H, T.M, T.G, PMH and DLW conceived the idea of the presented work and gave final approval of the version to be published. Specific contributions: A.T wrote ethics approvals, recruited subjects. PMH and DLW funded clinical chemistry. All authors took part in study design. A.T and DLW performed clinical chemistry and analyzed data. A.T and T.G performed all statistics and produced tables and figures. A.T and DLW worked on initial draft. All authors took part in finalizing the paper.

